# Structural insights into the activation of autoinhibited human lipid flippase ATP8B1 upon substrate binding

**DOI:** 10.1101/2021.11.07.467649

**Authors:** Meng-Ting Cheng, Yu Chen, Zhi-Peng Chen, Cong-Zhao Zhou, Wen-Tao Hou, Yuxing Chen

**Author notes:** Correspondence author: Yuxing Chen, Wen-Tao Hou, Cong-Zhao Zhou, **Email:** (Y.C.), (W.T.H) or (C.Z.Z.). **Author Contributions:** These authors contributed equally to this work (Meng-Ting Cheng, Yu Chen).

## Abstract

The human P4-type ATPase ATP8B1 in complex with the auxiliary noncatalytic protein CDC50A or CDC50B mediates the transport of cell membrane lipids from the outer to the inner membrane leaflet, which is crucial to maintain the asymmetry of membrane lipid. Its dysfunction usually leads to imbalance of bile acid circulation, and eventually causing intrahepatic cholestasis diseases. Here we found that both ATP8B1-CDC50A and ATP8B1-CDC50B possess a higher ATPase activity in the presence of the most favored substrate phosphatidylserine (PS); and moreover, the PS-stimulated activity could be augmented upon the addition of bile acids. The cryo-electron microscopy structures of ATP8B1-CDC50A at 3.36 Å and ATP8B1-CDC50B at 3.39 Å enabled us to capture an unprecedented phosphorylated and autoinhibited state, with the N- and C-terminal tails separately inserting into the cytoplasmic inter-domain clefts of ATP8B1. The PS-bound ATP8B1-CDC50A structure at 3.98 Å indicated the autoinhibited state could be released upon PS binding. Structural analysis combined with mutagenesis revealed the residues that determine the substrate specificity, and a unique positively charged loop in the phosphorylated domain of ATP8B1 for the recruitment of bile acids. Altogether, we updated the Post-Albers transport cycle, with an extra autoinhibited state of ATP8B1, which could be activated upon substrate binding. These findings not only provide structural insights into the ATP8B1-mediated restoration of human membrane lipid asymmetry during bile acid circulation, but also advance our understanding on the molecular mechanism of P-type ATPases.

## Introduction

The eukaryotic cell membrane consists of a variety of lipids, which are asymmetrically distributed across the lipid bilayer (1). Phosphatidylcholine (PC), sphingomyelin (SM) and glycolipids are enriched at the outer leaflet of the plasma membrane, whereas phosphatidylserine (PS), phosphatidylethanolamine (PE) and phosphatidylinositol (PI) are mainly restricted to the inner leaflet. The asymmetric distribution of lipids is essential to maintain cellular functions, including cell and organelle shape determination and dynamics, vesicle budding and trafficking, membrane stability and impermeability, cell signaling, apoptosis, and homeostasis of bile and cholesterol (2, 3). The disturbance of lipid asymmetry will lead to the imbalance of cell membrane and eventually cell death. For instance, loss of PS asymmetry is an early indicator of cell apoptosis, as well as a signal to initiate blood clotting (4). To maintain membrane asymmetry, eukaryotic cells express a series of cooperatively functioning lipid transporters, such as scramblases, floppases and flippases. Scramblases, which are energy independent, drive bidirectional lipid scrambling in response to intracellular concentrations of Ca^2+^ (5); whereas floppases, which are usually ATP-binding cassette (ABC) transporters, mediate the unidirectional translocation of lipids from the inner to the outer leaflet of the membrane bilayer (6). In contrast, flippases are usually type 4 P-type ATPase (P4-ATPase) in eukaryotic cells, which transport phospholipids from the outer leaflet to the inner leaflet (3).

The widespread P-type ATPases, which are featured with a phosphorylated intermediate during the transport cycle (P stands for phosphorylation), catalyze the transport of ions or phospholipids by utilizing the energy of ATP hydrolysis (7, 8). The human genome encodes a total of 14 P4-ATPase members, which are grouped into 5 classes, namely, Classes 1a, 1b, 2, 5 and 6 (3). For proper localization and integral function, P4-ATPases, except for Class 2 members (9), should form a complex with an auxiliary noncatalytic protein of the CDC50 family (10). To date, three CDC50 paralogs (CDC50A, CDC50B and CDC50C) have been identified in mammals (11). P4-ATPases from Class 1a, Class 5 and Class 6 bind to only CDC50A (10, 12), whereas those from Class 1b are able to form complexes with either CDC50A or CDC50B (12, 13). Recently, a series of structures of human P4-ATPases from Class 1a (ATP8A1) and Class 6 (ATP11C) complexed with CDC50A have been reported (14–16). However, the complex structure of P4-ATPase from Class 1b with CDC50A/B remains unknown.

ATP8B1, a member of Class 1b P4-ATPase, is also known as FIC1 (familial intrahepatic cholestasis type 1) (17). It colocalizes with the primary bile salt export pump ABCB11 in cholangiocytes and the canalicular membrane of hepatocytes (18). To counteract the disturbance of the asymmetric homeostasis of the cell membrane resulting from lipid flow accompanied by bile acid transport driven by ABCB11 (19), the floppase ABCB4 exports PC to envelop bile acid micelles (20), and ATP8B1 flips PS (21) or PC (22) from the outer to the inner membrane leaflet to restore membrane asymmetry. Defects in ATP8B1 are usually associated with severe human diseases, such as the intrahepatic cholestasis diseases PFIC1 (progressive familial intrahepatic cholestasis type 1) and BRIC1 (benign recurrent intrahepatic cholestasis type 1) (23).

Here, we solved the cryo-electron microscopy (cryo-EM) structures of apo-form ATP8B1-CDC50A and ATP8B1-CDC50B, and a PS-bound structure of ATP8B1-CDC50A. These structures unraveled a yet-unknown autoinhibited phosphorylated state of P-type ATPases, which is activated upon substrate binding. These findings not only advance our understanding of the molecular mechanism of P-type ATPases, but also provide a structural platform for further therapeutic intervention of intrahepatic cholestasis diseases.

## Results

### The ATPase activities of ATP8B1-CDC50A/B are significantly stimulated by PS

We overexpressed human ATP8B1-CDC50A and ATP8B1-CDC50B in HEK29F cells, with a 3×Flag tag fused to the N-terminus of ATP8B1, and purified the complexes using an affinity column followed by size-exclusion chromatography. SDS-PAGE indicated that both complexes were purified in a homogeneous state (Fig. S1). As substrates were reported to greatly enhance the ATPase activity of P4-ATPases (14, 16, 24), we performed ATPase activity in the presence or absence of different phospholipids at 300 μM. DDM (1%) was added as the hydrotropic agent for the dissolution of lipids to produce clear and homogeneous stock solutions at room temperature. The ATPase activities of both ATP8B1-CDC50A and ATP8B1-CDC50B showed a significant increase when PS was added compared to the basal activity (Fig. 1*A*). The *K_m_* values of PS-dependent ATPase activities for the ATP8B1-CDC50A and ATP8B1-CDC50B complexes were 92.6 ± 19.2 and 87.6 ± 9.1 μM, with *V_max_* values of 248.1 ± 22.3 and 195.4 ± 7.1 nmol min^-1^ mg^-1^, respectively (Fig. 1*B*). However, PC only showed a modest stimulation of the ATPase activity for ATP8B1-CDC50B but no significant effect for ATP8B1-CDC50A. In contrast, PE had no effect on ATPase activity stimulation for both complexes. These results revealed that both ATP8B1-CDC50A and ATP8B1-CDC50B possess a higher ATPase activity in the presence of PS, compared to PC.

**Figure 1.**
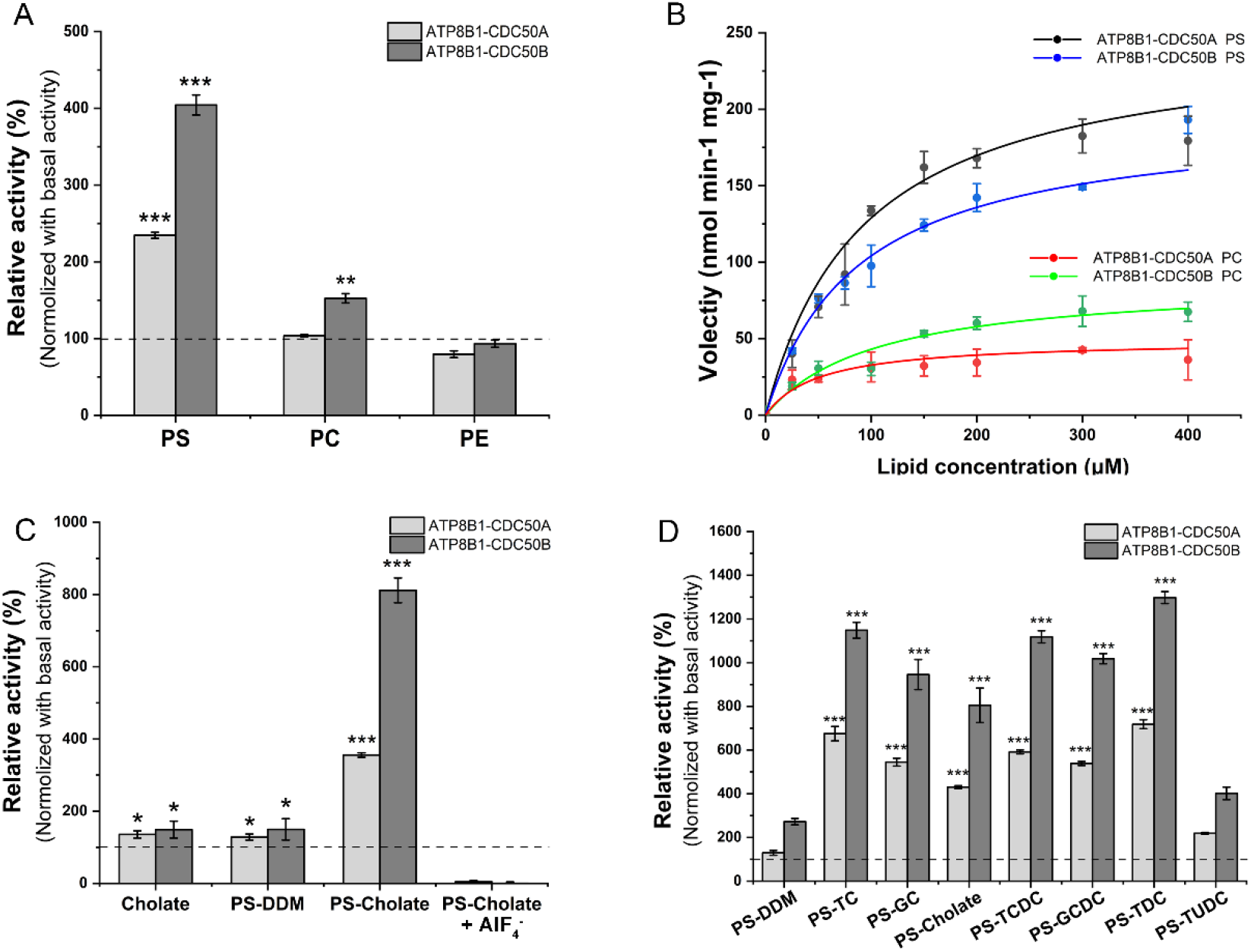
Substrate-stimulated and bile-acid-augmented ATPase activity assays of ATP8B1-CDC50A and ATP8B1-CDC50B complexes. (A) ATPase activities of ATP8B1-CDC50A/B in different phospholipids compared to the corresponding basal activities. The final concentration of the phospholipid was set to 300 μM. (B) The phospholipid concentration-dependent ATPase activities of ATP8B1-CDC50A/B. Data points represent the mean ± S.D. of three measurements at 37°C and were nonlinear fitted by the Michaelis-Menten equation. The *V_max_* values were calculated by the phosphate produced in mole per mg of ATP8B1-CDC50A or ATP8B1-CDC50B protein per min. (C) Cholate-augmented ATPase activities of ATP8B1-CDC50A/B in the presence of PS. The phospholipid was dissolved in DDM or cholate sodium at a final concentration of 100 μM. AlF_4_^-^ was produced by mixing AlCl3 and NaF. (D) Augmented ATPase activity of ATP8B1-CDC50A/B upon addition of PS dissolved in various bile acids. Abbreviations: TC, taurocholic acid; GC, glycocholic acid; TCDC, taurochenodoxycholic acid; GCDC, glycochenodeoxycholic acid; TDC, taurodeoxycholic acid; TUDC, tauroursodeoxycholate acid. Data were normalized against the basal activities of ATP8B1-CDC50A or ATP8B1-CDC50B, respectively. At least three independent assays were performed to calculate the means and standard deviations, and the data are presented as the means ± S.D. Two-tailed Student’s t-test was used for the comparison of statistical significance. The p values of <0.05, 0.01 and 0.001 are indicated with *, ** and ***, respectively.

### Bile acids can further augment the ATPase activity

It is puzzling that the *V_max_* of ATP8B1 is less than 1/6 of the previously reported P4-ATPases (14, 16, 24). Notably, the yeast P4-ATPase Drs2p in complex with Cdc50p with a non-detectable ATPase activity in the presence of substrate, could be activated by an activator molecule, lipid phosphatidylinositol 4-phosphate (PI4P) (25). In addition, bile acids could enhance the ATPase activities of ABCB4 (26) and ABCG5/G8 (27), both of which are localized in hepatocytes like ATP8B1. Thus, we tested the ATPase activity of ATP8B1-CDC50A/B complexes in the presence of cholate, the main component of bile acids. However, the addition of cholate alone only exhibited a slight increase in activity for both complexes, compared to the basal activity (Fig. 1*C*). Compared to PS solubilized in the detergent DDM with ~50% activity increase, the two complexes in the presence of PS solubilized in cholate displayed 4- and 8-fold activities, respectively (Fig. 1*C*). Notably, the activity was completely inhibited by AlF_4_^-^, an inhibitor of P-type ATPases (28). The ATPase activity assays in the presence of PS solubilized in other detergents, such as OG and GDN, gave a result similar to that in DDM (Fig. S2). Moreover, PS solubilized in either primary or secondary conjugated bile acids could greatly augment the ATPase activity (Fig. 1*D*), compared to that in DDM. Notably, tauro-conjugated bile acids, including taurocholic acid (TC), have higher augmentation rates. All these data suggested that various bile acids can augment the PS-simulated ATPase activity of ATP8B1-CDC50A/B.

### Overall structures of ATP8B1-CDC50A and ATP8B1-CDC50B

We solved the cryo-EM structures of the ATP8B1-CDC50A and ATP8B1-CDC50B complexes at 3.36 and 3.39 Å, respectively (Fig. 2*A* and Fig. 2*B*), which followed the gold-standard Fourier shell correlation criterion of 0.143 (Fig. S3 and Fig. S4). Superposition of the two complex structures yields a root-mean-square deviation (RMSD) of 0.42 Å over 990 Cα atoms, indicating that the two structures resemble each other. Similar to the previously reported P4-ATPase structures, ATP8B1 also has a TM (transmembrane) domain of 10 TMs (TM1-10) and three cytoplasmic domains: the A (actuator), N (nucleotide) and P (phosphorylation) domains (Fig. 2*A* and Fig. 2*B*). The N domain is responsible for binding to ATP, which donates the phosphate group for autophosphorylation of the conserved Asp454 of the P domain, generating a phosphorylated intermediate during the transport cycle. The A domain can dephosphorylate the phosphorylated P domain via the catalytic residue Glu234, and finally triggering substrate translocation. In addition, the N-terminus and C-terminus form long loops, termed the N-tail and C-tail, respectively, which might function as regulatory domains (29). In both complexes, superposition of CDC50A and CDC50B gives a quite similar structure with an RMSD of 0.79 Å over 269 Cα atoms, in agreement with their high sequence identity of 54% (Fig. S5). CDC50A/B interact with ATP8B1 through the extracellular domains, the two TMs and an unstructured loop proceeding the N-terminus of the TMs, forming three interfaces almost identical in the two complexes. These interfaces are conserved in known structures of P4-ATPases complexed with CDC50A (14, 15).

**Figure 2.**
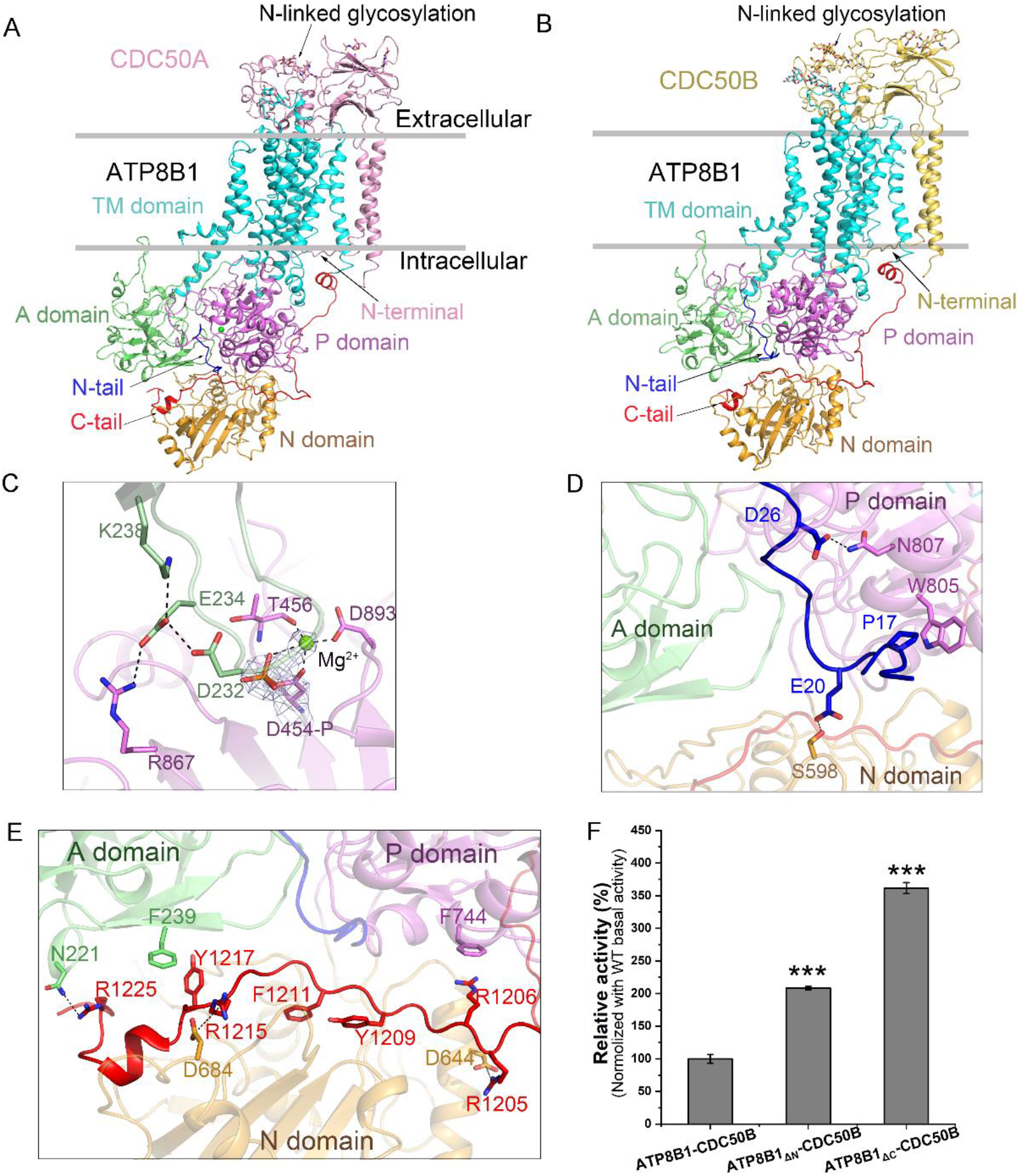
Structures of ATP8B1-CDC50A and ATP8B1-CDC50B at the autoinhibited phosphorylated state. Overall structure<u>s</u> of (A) ATP8B1-CDC50A and (B) ATP8B1–CDC50B in cartoon. The major domains and motifs are labeled in different colors. The same color scheme is used throughout the manuscript. (C) The phosphorylation site. The phosphorylated Asp454 from P domain and related residues are shown in sticks. PO4^-^ is shown in sticks, and Mg^2+^ is shown as spheres. Densities are shown as gray mesh, contoured at 8σ. Interactions of (D) the N-tail and (E) the C-tail with the cytoplasmic domains. All interacting residues are shown in sticks. (F) ATPase activities of the N-tail or C-tail truncated ATP8B1 in complex with CDC50B compared to the wild type. △N and △C stand for the deletion of N-terminal Met1~Glu30 and C-terminal Ala1187~Ser1251, respectively. Data were normalized against the basal activity of ATP8B1-CDC50B. At least three independent assays were performed to calculate the means and standard deviations, and the data are presented as the means ± S.D. Two-tailed Student’s t-test was used for the comparison of statistical significance. The p values of <0.05, 0.01 and 0.001 are indicated with *, ** and ***, respectively.

### ATP8B1 adopts an autoinhibited E2P state

The transport cycle of P4-ATPases has been clearly depicted by the Post-Albers scheme (30, 31), including several intermediate states (Fig. S6): ATP binding (E1-ATP), phosphorylated (E1P and E2P), substrate binding (E2Pi-PL), dephosphorylated (E2) and substrate release (E1) state (8). These states have been captured in a previous report of the cryo-EM structure of ATP8A1-CDC50A (14). Superposition of ATP8B1 with the known structures of ATP8A1 revealed that ATP8B1 possesses a lowest RMSD of 2.01 Å with ATPB8A1 in the E2P state (Fig. S7).

In our two structures, an extra density could be found at the proximity of residue Asp454 from the conserved DKGT motif of the P domain, which could be fitted with a PO_4_^-^ and a Mg^2+^ ion (Fig. 2*C*), indicating that Asp454 is phosphorylated. The Mg^2+^ ion is coordinated by Asp454, Thr456, Asp893 and PO_4_^-^ (Fig. 2*C*). However, the catalytic residue Glu234 from the conserved DGET motif of the A domain is stabilized by Asp232, Lys238 and Arg867 via salt bridges and hydrogen bonds. As a result, the phosphorylated Asp454 is too far away from Glu234 to be dephosphorylated, which is a feature of the E2P state (14). Thus, ATP8B1 is most likely in the E2P state, in which the P domain has been phosphorylated and ready to bind the substrate.

Of note, both the N-tail and C-tail of ATP8B1 insert into the clefts of the three cytoplasmic domains (Fig. 2*A* and Fig. 2*B*). Specifically, the N-tail interacts with the P domain by two pairs of hydrogen bonds (Glu20-Ser598 and Asp26-Asn807) and hydrophobic interactions between Pro17 and Trp805 (Fig. 2*D*). The C-tail interacts with the P domain via the cation-π interaction between Arg1206-Phe744, the A domain via π-π interaction between Phe239-Tyr1217 and a hydrogen bond between Asn221-Arg1225, and the N domain via two pairs of salt bridges between Asp644-Arg1205 and Asp684-Arg1215 (Fig. 2*E*). Remarkably, Phe1211 from the conserved GYAFS motif occupies the ATP binding site (Fig. 2*E*). Truncation of either the N-tail or C-tail of ABT8B1 resulted in an increased ATPase activity of ~2 or 3-fold to that of the wild type, respectively (Fig. 2*F*), further suggesting that ATP8B1 basically adopts an autoinhibited E2P state.

### The positively charged P-loop is responsible for the recruitment of bile acids

Structural analysis revealed three positively charged regions on ATP8B1: an outstretched P-loop from the P domain (residues Lys813-Lys846) and two segments from the C-tail, which are termed C-helix (residues Ser1172-Arg1184) and C-turn (residues Arg1193-Arg1206), respectively (Fig. 3*A*). There are 16 positively charged residues in the 34-residue P-loop of ATP8B1, which is absent in other P4-type ATPases (Fig. 3*B*). C-helix is a short helix corresponding to the amphipathic helix of yeast Drs2p, which is adjacent to the binding site of the activator PI4P (32), while C-turn is an arginine-rich sharp turn of C-tail (Fig. S8).

**Figure 3.**
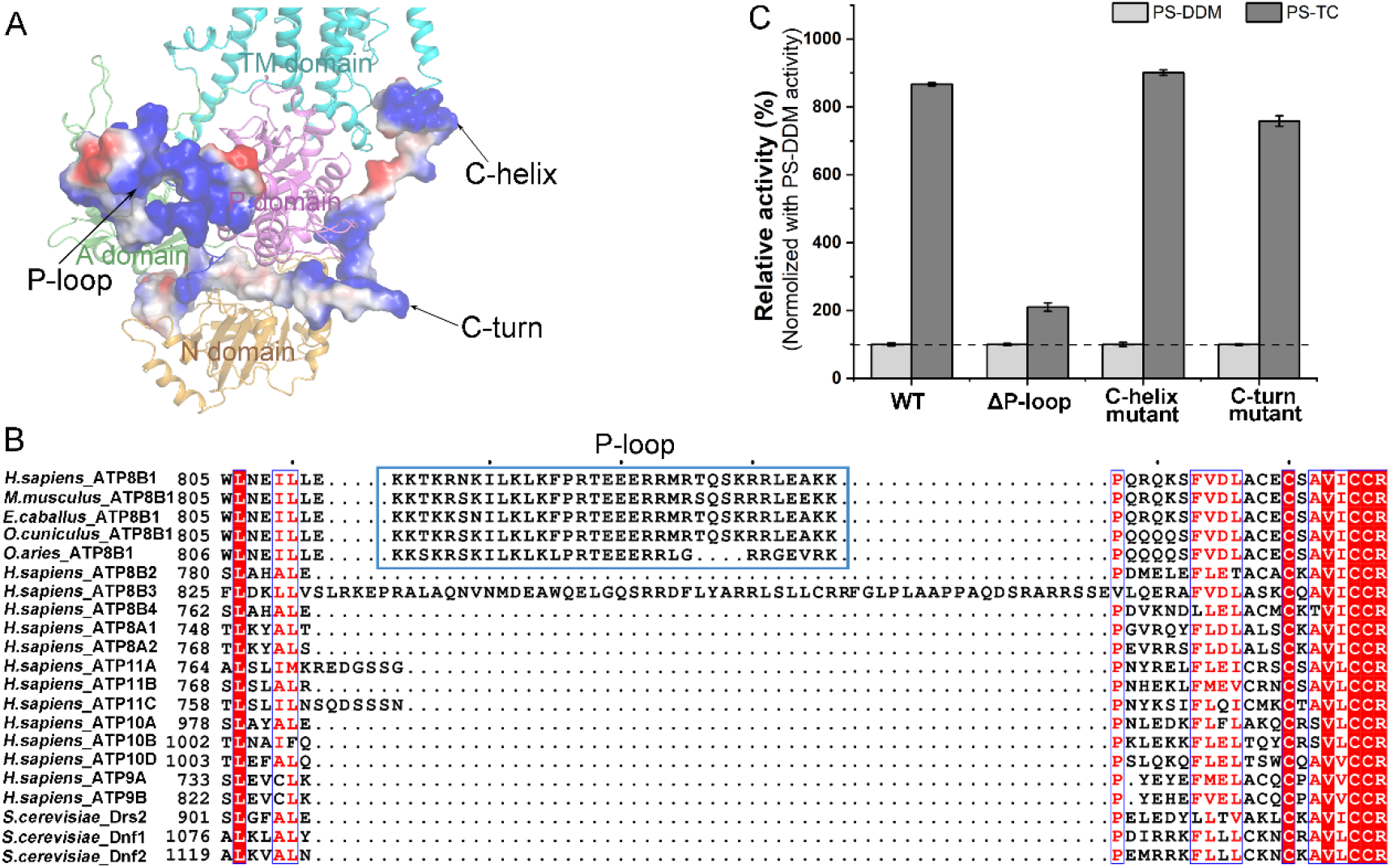
The P-loop conserved in ATP8B1 and homologs is essential for the recruitment of bile acids. (A) The three positively charged regions of ATP8B1, namely the P-loop, C-helix and C-turn. (B) Multiple-sequence alignment of the P-loop and flanking regions in P4-type ATPases. The P-loop rich of positively charged residues in ATP8B1 and homologs are marked with a blue box. (C) ATPase activities of ATP8B1-CDC50B mutants compared to the wild type, in the presence of 100 μM PS dissolved in either DDM or TC. The activities in the presence of PS dissolved in TC were normalized against the corresponding activity in the presence of PS dissolved in DDM. The ΔP-loop means deletion of K813-K846 of ATP8B1, whereas C-helix and C-turn mutants represent the multiple mutations of K1177E-K1180E-H1181D-R1182E-K1183E-R1184E-K1186E and R1194T-R1199S-R1200S-R1206S, respectively. At least three independent assays were performed to calculate the means and standard deviations, and the data are presented as the means ± S.D.

Hence, we truncated the P-loop (residues from Lys813 to Lys846) and mutated the positively charged residues in C-helix (K1177E-K1180E-H1181D-R1182E-K1183E-R1184E-K1186E) and C-turn (R1194T-R1199S-R1200S-R1206S) for ATPase activity assays. The results showed that the truncation of P-loop led to a significant reduction in TC-augmented activity compared to the wild type, whereas either C-helix or C-turn mutant displayed no change in activity (Fig. 3*C*). It suggested that the P-loop, which is conserved in ATP8B1 homologs according to the multisequence alignment (Fig. 3*B*), is responsible for the recruitment of bile acids.

### The substrate-bound ATP8B1-CDC50A structure revealed the release of autoinhibition

To further investigate the substrate specificity of ATP8B1, we solved the cryo-EM structure of substrate-bound ATP8B1-CDC50A at a 3.98 Å resolution (Fig. S9). An extra density in the cavity formed by TM2, TM4 and TM6 of ATP8B1 could be fitted with a molecule of PS (Fig. 4*A*). The serine group of PS is stabilized by Thr143, Thr144 and Asn397, which are conserved in PS flippases (Fig. S10). The phosphate group and glycerol backbone of PS are embraced by Ser403, Asn989 and Ser994 (Fig. 4*B*), which are conserved residues in most P4-ATPases (Fig. S10). Notably, previous reports showed that clinical variants of S403Y and S994R of ATP8B1 were associated with the severe intrahepatic cholestasis disease PFIC1 (33, 34). Moreover, single mutation (T143A, T144A, N397A, S403Y, N989A or S994R) of the PS binding residues showed a significantly reduced ATPase activity in the presence of PS dissolved in TC (Fig. 4*C*).

**Figure 4.**
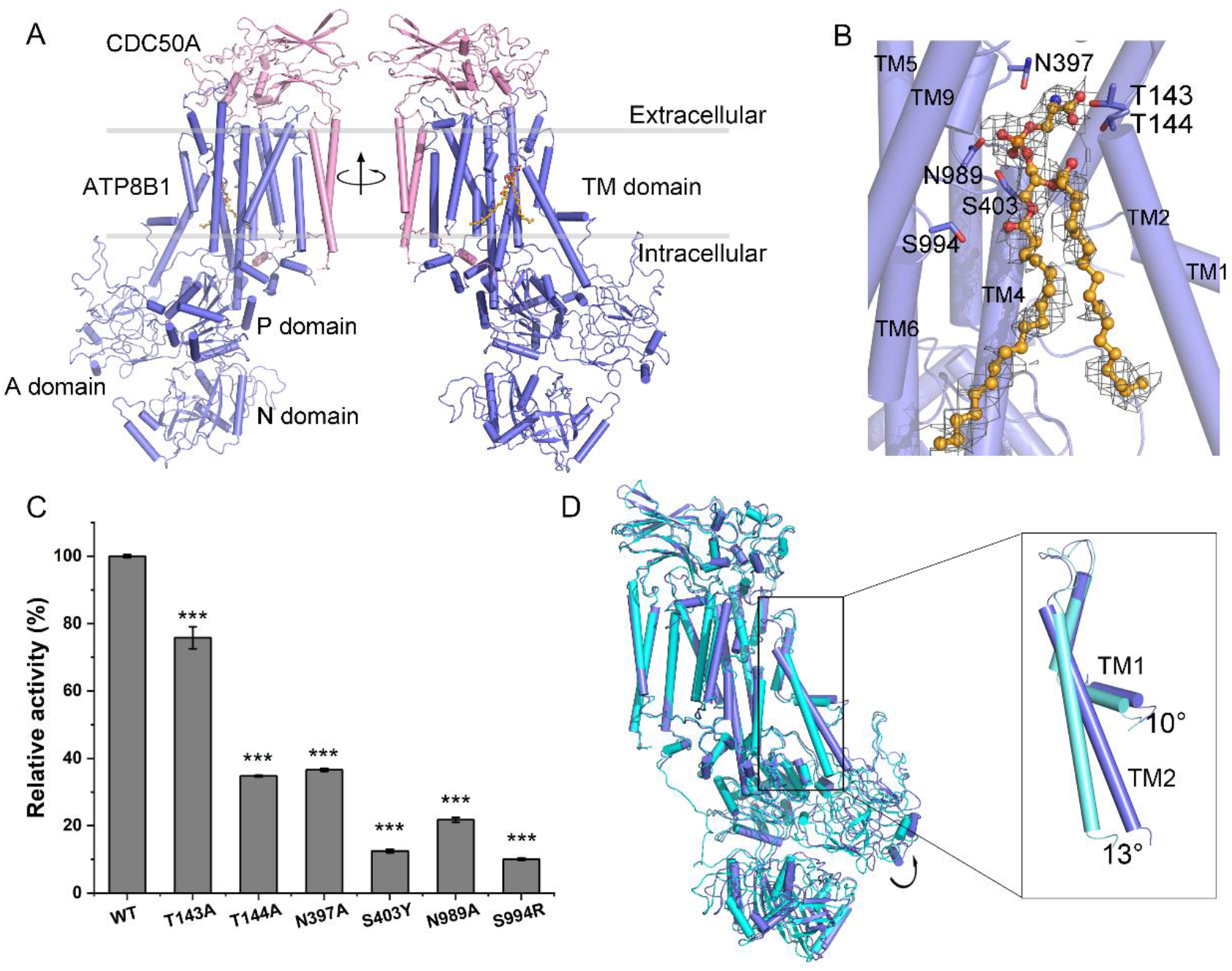
The PS-bound structure of ATP8B1-CDC50A and the PS-binding site. (A) Overall structure of PS-bound ATP8B1-CDC50A. ATP8B1 is colored in slate, PS is shown in sticks and spheres in orange. (B) The PS-binding site. The density of the PS is shown in gray mesh, contoured at 3σ. (C) ATPase activities of ATP8B1-CDC50B mutants compared to the wild type, in the presence of 100 μM PS dissolved in TC. Data were normalized against activity of the wild type. At least three independent assays were performed to calculate the means and standard deviations, and the data are presented as the means ± S.D. Two-tailed Student’s t-test is used for the comparison of statistical significance. The p values of <0.05, 0.01 and 0.001 are indicated with *, ** and ***, respectively. (D) Superposition of autoinhibited E2P state and PS-bound state of ATP8B1-CDC50A by aligning the TM7-10 of ATP8B1. The E2P state ATP8B1 is shown in cyan.

Remarkably, both the N-tail and C-tail of ATP8B1 are invisible in the PS-bound structure (Fig. 4*A*), suggesting the release of autoinhibition upon substrate binding. Superposition of this PS-bound structure against the apo-form ATP8B1 revealed significant rotations of TM1 and TM2 for ~10° and ~13°, respectively. In consequence, the A domain shifts towards the P domain, which is subject to being dephosphorylated (Fig. 4*D*). This conformation is similar to that of ATP8A1 in the E2Pi-PL state (14), in which AlF_4_^-^ substituted PO4^-^ at the proximity of residue Asp454 of the P domain, despite we could not fit AlF_4_^-^ in our PS-bound structure due to the relatively low resolution of cytoplasmic domains (Fig. S9).

## Discussion

An autoinhibited state has been reported in the previous structures of yeast Drs2p, the C-tail of which was found to lock the cytoplasmic domains (32, 35). In addition, human ATP8A1 can be locked in an inhibited E2P state upon addition of the inhibitor BeF3^-^ (14). Our present structures, in complex with either CD50A or CDC50B, showed that ATP8B1 adopts an autoinhibited E2P state, in which the N-tail and C-tail separately insert into the cytoplasmic inter-domain clefts and the residue Asp454 of the P domain is phosphorylated. ATPase activity assays indicated that the activity of ATPB1 could be inhibited by its own N-tail and C-tail, which might interrupt the inter-domain crosstalk. Most likely, an inhibited E2P state for inhibiting the activity of P-type ATPases is necessary at the homeostasis of membrane lipid asymmetry.

Based on our findings and previous reports, we updated the Post-Albers cycle with an extra autoinhibited E2P state, which is an equilibrium state with the E2P state (Fig. 5). As shown in our PS-bound structure, this autoinhibition state could be switched to the E2Pi-PL state upon substrate binding (Fig. 5). In addition, our biochemical results indicated that bile acids can enhance the PS-stimulated ATPase activity of ATP8B1, similar to the previous reports that PI4P can stimulate the ATPase activity of yeast Drs2p (25, 36). Furthermore, deletion of the positively charged P-loop, which is highly conserved in ATP8B1 homologs, led to a reduced ATPase activity of ATP8B1 in response to the addition of bile acids. In fact, ATP8B1 usually functions under a physiological environment with enriched bile acids; thus, the release of autoinhibition could be induced by the substrate, and further accelerated upon the easily recruited bile acids.

**Figure 5.**
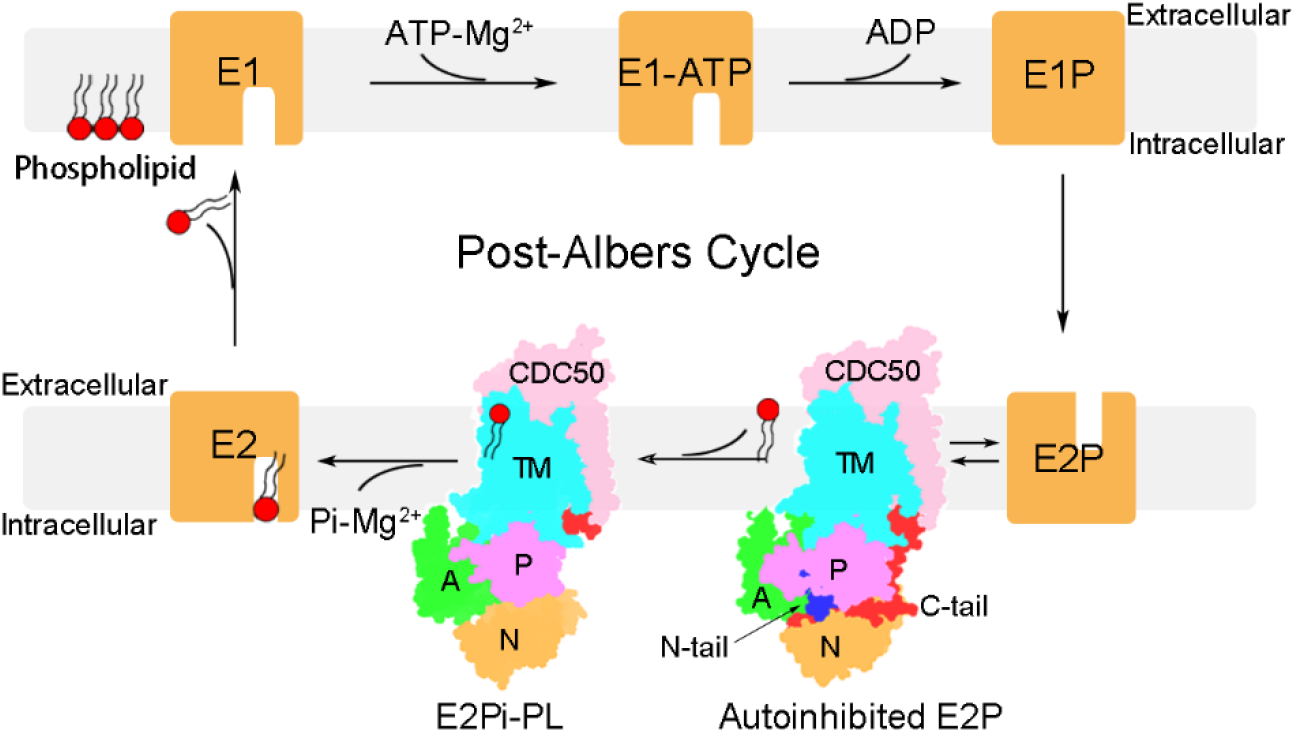
An updated model of Post-Albers cycle. The two structures (autoinhibited E2P and E2Pi-PL) we report here are shown as the schemes, with the domains of ATP8B1-CDC50 shown in different colors, whereas the rest states are shown as yellow squares. Binding and hydrolyzing ATP would trigger transition of ATP8B1 from the state E1 to E1P. Afterwards, the rearrangement of the cytoplasmic domains makes ATP8B1 to adopt an E2P state, which is most likely equilibrated at the autoinhibited E2P state with the N-tail and C-tail inserting into the cytoplasmic inter-domain clefts. The release of autoinhibition could be induced upon phospholipid binding and further accelerated in the presence of bile acids, switching ATP8B1 at the E2Pi-PL state. The ATP8B1 at the E2 state with a dephosphorylated P domain undergoes domain rearrangement accompanying with the release of phospholipid; and finally ATP8B1 returns to the E1 state.

In summary, ATP8B1 free of substrate basically adopts an autoinhibited conformation at the homeostatic membrane asymmetry. Once this asymmetry is altered, usually due to the phospholipid flow accompanying with the efflux of bile acids across the membrane, ATP8B1 is fully activated in the presence of both substrate and bile acids. The present structural analysis together with biochemical assays updated our understanding on the Post-Albers cycle.

## Materials and Methods

### Cloning and expression

The full-length human *ATP8B1* (Uniprot: O43520), *CDC50A* (Uniprot: Q9NV96) and *CDC50B* (Uniprot: Q3MIR4) genes were synthesized after codon optimization for the mammalian cell expression system by Sangon Biotech Company. The wild-type *ATP8B1* and mutants were subcloned into a pCAG vector with an N-terminal Flag-tag (DYKDDDDK). *CDC50A* and *CDC50B* were respectively subcloned into the same vector with a C-terminal 6×His-tag or an N-terminal 6×His-tag, using One Step Cloning Kit (Vazyme).

For protein expression, HEK293F cells were cultured in SMM 293T-II medium (Sino Biological Inc.) at 37°C with 5% CO2. Cells were transfected when the density reached ~2.5 × 10^6^ cells per mL. For transfection, ~1.8 mg pCAG-ATP8B1 and ~0.3 mg pCAG-CDC50A/B were incubated with 4 mg linear polyethylenimines (PEIs) (Polysciences, Inc) in 45 mL fresh medium for 15 min, followed by a 15-min static incubation. The transfected cells were grown at 37°C with 5% CO2 for 48 h before harvesting. Cell pellets were resuspended in the lysis buffer containing 50 mM Tris-HCl pH 7.5, 150 mM NaCl, 20% glycerol (w/v) after centrifugation at 1,500 g for 10 min. The suspension was frozen in liquid nitrogen and stored at −80°C for further use.

All mutants were generated with a standard PCR-based strategy and were cloned, overexpressed and purified the same way as the wild-type protein.

### Protein preparation

For protein purification, 2 mM ATP (Sangon) and 2 mM MgCl2 were added to the thawed suspension and the mixture was incubated with additional 1% (w/v) dodecyl-β-D-maltopyranoside (DDM; Bluepus) and 0.2% (w/v) cholesteryl hemisuccinate (CHS, Anatrace) at 8°C for 2 h for membrane solubilization and protein extraction. After ultracentrifugation at 45,000 rpm for 45 min (Beckman Type 70 Ti), the supernatant was incubated with the anti-Flag M2 affinity gel (Sigma) on ice for at least 40 min. Then the resin was washed by 30 mL of wash buffer containing 50 mM Tris-HCl pH 7.5, 150 mM NaCl, 10% (w/v) glycerol, with 0.02% glyco-diogenin (GDN, w/v, Anatrace) for ATP8B1-CDC50A or 0.06% digitonin (w/v, Apollo Scientific) for ATP8B1-CDC50B. The protein was eluted with 6 mL of wash buffer plus 200 μg/ml Flag peptide. The samples were further concentrated and purified by size-exclusion chromatography (SEC) on a Superose 6 Increase 10/300 GL column (GE Healthcare), pre-equilibrated with SEC buffer (50 mM Tris-HCl pH 7.5, 150 mM NaCl, and 0.02% GDN or 0.06% digitonin). The peak fractions were collected, concentrated to 4-6 mg/mL, frozen in liquid nitrogen and stored at −80°C before use.

### Lipid and detergent/lipid mixture preparation

Lipids were prepared in 20 mM Tris-HCl pH 7.5, 75 mM KCl to a final concentration 10 mg/mL (~12.5 mM) by sonicating in a water bath sonicator at room temperature until forming a uniform solution. Bile acids (sodium salts) were dissolved at a final concentration of 100 mM in water and mixed with lipid stocks in a mole ratio of 3:1. 20% (w/v) DDM stocks were prepared in water and added to lipids solution in a final concentration of 1%. All the mixtures were frozen in liquid nitrogen and thawed at room temperature for 3 times, resulting in a clear and homogeneous solution at room temperature. Lipid stocks, bile acids and lipid mixtures were stored at −20°C before use.

### ATPase activity assays

For the substrate-simulated ATPase activity assay, 0.01 mg (~0.05 μM) of the purified proteins were pre-incubated at room temperature for 5 min with or without 600 μM lipids in 150 μL reaction buffer, containing 20 mM Tris-HCl pH 7.5, 75 mM KCl, 2 mM MgCl_2_, 0.02% DDM. The proteins or protein/lipid mixtures were cooled in ice and further mixed with an equal volume of pre-cooled reaction buffer containing 4 mM ATP. Reactions were performed at 37°C for 20 min and the amount of released phosphate group (Pi) was quantitatively measured using the ATPase colorimetric Assay Kit (Innova Biosciences) in 96-well plates at OD_650_ nm.

The bile-acid-augmented assays were measured similar to that mentioned above, except that the proteins were pre-incubated with detergent/lipids mixtures to a final lipid concentration of 200 μM.

### Cryo-EM sample preparation

For the apo-form ATP8B1-CDC50A or ATP8B1-CDC50B complex, purified proteins were concentrated to 4.5 mg/mL. After centrifugation at 12,000 rpm for 10 min, 3.5 μL samples were placed on glow-discharged holey carbon grids (Quantifoil, Cu R1.2/1.3, 300-mesh) with the force −2, blot time 3 sec and plunged into liquid ethane by using Vitrobot Mark IV (FEI) at 8°C and 100% humidity.

For the substrate binding sample, ATP8B1-CDC50A purified by the anti-Flag M2 affinity gel were pre-incubated with 2 mM AlCl3, 10 mM NaF and 2 mM MgCl_2_ overnight. The mixture samples were further concentrated and purified by SEC. The purified ATP8B1-CDC50A were concentrated to 5 mg/mL, incubated with additional 1 mM AlCl3, 5 mM NaF, 2 mM MgCl_2_ and 10 μM 1,2-dioleoyl-sn-glycero-3-phospho-L-serine (DOPS) for 1 h in ice. 3.5 μL samples were placed on glow-discharged holey carbon grids (Quantifoil, Cu R1.2/1.3, 300-mesh) with the force −2, blot time 3 sec and plunged into liquid ethane by using Vitrobot Mark IV (FEI) at 8°C and 100% humidity.

### Cryo-EM data collection

The cryo-EM grids of apo-form and PS-bound ATP8B1-CDC50A were loaded into a Titan Krios transmission electron microscope (ThermoFisher Scientific) operating at 300 KeV with a Gatan K2 Summit direct electron detector at the Center for Integrative Imaging of Hefei National Laboratory for Physical Sciences at the Microscale, University of Science and Technology of China (USTC). A total of 3003/2983 movie stacks were collected in super resolution mode at nominal magnification of 29,000 × with a defocus range from −2.5 to −1.5 μm. Each movie stack of 32 frames was exposed for 6.4 sec under a dose rate of 10 e/pixel/sec, resulting in a total dose of ~60 e Å^-2^.

The cryo-EM data of apo-form ATP8B1-CDC50B complex were collected at the Center for Biological Imaging at the Institute of Biophysics (IBP), Chinese Academy of Sciences (CAS). A total of 3717 micrographs were collected in super resolution mode with K3 camera at nominal magnification of 22,500 × with a defocus range from −1.5 to −2.0 μm. Exposures of 6.4 s fractionated into 32 frames were collected at a dose rate of 1.5 or 1.6 e^-^ per Å^2^ per frame, corresponding to a total dose of ~60 e-per Å^2^.

### Cryo-EM data processing

All movie frames were corrected for gain reference and binned by a factor of 2 to yield a pixel size of 1.06 Å in RELION3.1 (37) through MotionCor2 (38). The contrast transfer function (CTF) parameters were estimated from the corrected movie frames using CTFFIND4 (39). After manual inspection of the micrographs, approximately 3,000 particles were manually selected. Particles were automatically extracted by RELION with binning factor 2. For apo-form ATP8B1-CDC50A, a total of 773,161 particles were picked and subjected to 2D classification. After multi-rounds of 2D classification, 399,091 particles were selected for further 3D classification with 3 classes using the reference generated by the 3D initial model. 182,618 particles from the best class were refined and re-extracted for further 3D refinement. To improve the EM density, 3D skip alignment classification, followed by CTF refinement and Bayesian Polishing were performed, giving rise to an average resolution of 3.36 Å.

For apo-form ATP8B1-CDC50B, 2,561,714 particles were automatically extracted and subjected to 2D classification. 494,609 particles were selected for further 3D classification with 4 classes using the reference generated by the 3D initial model. 160,435 particles from one of the classes were further refined and post-processed to yield a 3.39 Å map.

For PS-bound ATP8B1-CDC50A, 1,286,936 particles were automatically extracted and subjected to 2D classification. 159,960 particles were selected for further 3D classification with 3 classes using the reference generated by the 3D initial model. 159,960particles from the best class were refined and re-extracted for further 3D refinement. 3D skip alignment classification, followed by CTF refinement and Bayesian Polishing were performed, giving rise to an average resolution of 3.98 Å.

The data processing pipelines are presented in SI Appendix. Map resolution was estimated with the gold-standard Fourier shell correlation 0.143 criterion (40). Local resolutions were estimated using Resmap with RELION3.1.

### Model building and refinement

The final sharpened map with a B-factor of −140 Å^2^ was used for model building in Coot (41). Initial structure models for ATP8B1 and CDC50B were predicted by SWISS-MODEL (42). The CDC50A structure was obtained from PDB 6K7L. The initial model of ATP8B1-CDC50A/B complexes were built by fitting the ATP8B1 and CDC50A/B model into the map using the UCSF Chimera. Then model building and refinement was accomplished manually by Coot (41). After several rounds of manual building, the model was almost completed built and automatically refined against the map by the real_space_refine program in PHENIX (43) with secondary structure and geometry restraints. Figures were prepared with Pymol (44) or Chimera (45).

## Supporting information

Figures S1 to S10;Table S1;SI References

## Acknowledgments

We thank Dr. Yongxiang Gao at the Center for Integrative Imaging, Hefei National Laboratory for Physical Sciences at the Microscale, University of Science and Technology of China, Xu-jing Li and the collogues at the Center for Biological Imaging at the Institute of Biophysics (IBP), Chinese Academy of Sciences (CAS) for the cryo-EM image acquisition. This work was supported by the Ministry of Science and Technology of China (2019YFA0508500), the Strategic Priority Research Program of the Chinese Academy of Sciences (XDB37020202).

## Data availability

The cryo-EM structures and cryo-EM density maps of the autoinhibited E2P state ATP8B1-CDC50A, the autoinhibited E2P state ATP8B1-CDC50B and PS-bound ATP8B1-CDC50A have been respectively deposited at PDB under the accession code of 7VGI, EMD-31970; EMD-31969, 7VGH; 7VGJ, EMD-31971.

## Author contributions

Yuxing Chen and Wen-Tao Hou conceived, designed and supervised the project. Meng-Ting Cheng designed and performed the experiments. Zhi-Peng Chen and Yu Chen. collected the Cryo-EM data and Yu Chen solved the structure. Meng-Ting Cheng and Wen-Tao Hou analyzed the data. Meng-Ting Cheng, Wen-Tao Hou, Yuxing Chen. and Cong-Zhao Zhou prepared the manuscript. All authors discussed the data and read the manuscript.

## References

1. P. A. Leventis, S. Grinstein, The distribution and function of phosphatidylserine in cellular membranes. Annu Rev Biophys 39, 407–427 (2010).

2. J. C. Holthuis, A. K. Menon, Lipid landscapes and pipelines in membrane homeostasis. Nature 510, 48–57 (2014).

3. J. P. Andersen et al., P4-ATPases as Phospholipid Flippases-Structure, Function, and Enigmas. Front Physiol 7, 275 (2016).

4. E. M. Bevers, P. L. Williamson, Getting to the Outer Leaflet: Physiology of Phosphatidylserine Exposure at the Plasma Membrane. Physiol Rev 96, 605–645 (2016).

5. H. M. Hankins, R. D. Baldridge, P. Xu, T. R. Graham, Role of flippases, scramblases and transfer proteins in phosphatidylserine subcellular distribution. Traffic 16, 35–47 (2015).

6. M. M. Babashamsi, S. Z. Koukhaloo, S. Halalkhor, A. Salimi, M. Babashamsi, ABCA1 and metabolic syndrome; a review of the ABCA1 role in HDL-VLDL production, insulin-glucose homeostasis, inflammation and obesity. Diabetes Metab Syndr 13, 1529–1534 (2019).

7. M. Dyla, M. Kjaergaard, H. Poulsen, P. Nissen, Structure and Mechanism of P-Type ATPase Ion Pumps. Annu Rev Biochem 89, 583–603 (2020).

8. J. A. Lyons, M. Timcenko, T. Dieudonne, G. Lenoir, P. Nissen, P4-ATPases: how an old dog learnt new tricks - structure and mechanism of lipid flippases. Curr Opin Struct Biol 63, 65–73 (2020).

9. H. Takatsu et al., ATP9B, a P4-ATPase (a putative aminophospholipid translocase), localizes to the trans-Golgi network in a CDC50 protein-independent manner. J Biol Chem 286, 38159–38167 (2011).

10. K. Segawa, S. Kurata, S. Nagata, The CDC50A extracellular domain is required for forming a functional complex with and chaperoning phospholipid flippases to the plasma membrane. J Biol Chem 293, 2172–2182 (2018).

11. Y. Katoh, M. Katoh, Identification and characterization of CDC50A, CDC50B and CDC50C genes in silico. Oncol Rep. 12:, 939–943 (2004).

12. S. Bryde et al., CDC50 proteins are critical components of the human class-1 P4-ATPase transport machinery. J Biol Chem 285, 40562–40572 (2010).

13. L. M. van der Velden et al., Heteromeric interactions required for abundance and subcellular localization of human CDC50 proteins and class 1 P4-ATPases. J Biol Chem 285, 40088–40096 (2010).

14. K. Y. M. Hiraizumi, T. Nishizawa, O. Nureki, Cryo-EM structures capture the transport cycle of the P4-ATPase flippase Science 365, 1149–1155 (2019).

15. H. Nakanishi et al., Transport Cycle of Plasma Membrane Flippase ATP11C by Cryo-EM. Cell Rep 32, 108208 (2020).

16. H. Nakanishi et al., Crystal structure of a human plasma membrane phospholipid flippase. J Biol Chem 295, 10180–10194 (2020).

17. L. N. Bull et al., A gene encoding a P-type ATPase mutated in two forms of hereditary cholestasis. Nat Genet 18, 219–224 (1998).

18. E. F. Eppens et al., FIC1, the protein affected in two forms of hereditary cholestasis, is localized in the cholangiocyte and the canalicular membrane of the hepatocyte. J Hepatol 35, 436–443 (2001).

19. B. Stieger, The role of the sodium-taurocholate cotransporting polypeptide (NTCP) and of the bile salt export pump (BSEP) in physiology and pathophysiology of bile formation. Handb Exp Pharmacol 201, 205–259 (2011).

20. A. R. Crawford et al., Hepatic secretion of phospholipid vesicles in the mouse critically depends on mdr2 or MDR3 P-glycoprotein expression. Visualization by electron microscopy. J Clin Invest 100, 2562–2567 (1997).

21. C. C. Paulusma et al., ATP8B1 requires an accessory protein for endoplasmic reticulum exit and plasma membrane lipid flippase activity. Hepatology 47, 268–278 (2008).

22. H. Takatsu et al., Phospholipid flippase activities and substrate specificities of human type IV P-type ATPases localized to the plasma membrane. J Biol Chem 289, 33543–33556 (2014).

23. L. N. Bull, R. J. Thompson, Progressive Familial Intrahepatic Cholestasis. Clinics in Liver Disease 22, 657–669 (2018).

24. Y. He, J. Xu, X. Wu, L. Li, Structures of a P4-ATPase lipid flippase in lipid bilayers. Protein Cell 11, 458–463 (2020).

25. X. Zhou, T. T. Sebastian, T. R. Graham, Auto-inhibition of Drs2p, a Yeast Phospholipid Flippase, by Its Carboxyl-terminal Tail. J Biol Chem 288, 31807–31815 (2013).

26. T. Kroll, S. H. J. Smits, L. Schmitt, Monomeric bile acids modulate the ATPase activity of detergent-solubilized ABCB4/MDR3. J Lipid Res 62, 100087 (2021).

27. B. J. Johnson, J. Y. Lee, A. Pickert, I. L. Urbatsch, Bile acids stimulate ATP hydrolysis in the purified cholesterol transporter ABCG5/G8. Biochemistry 49, 3403–3411 (2010).

28. L. Bai et al., Transport mechanism of P4 ATPase phosphatidylcholine flippases. Elife 9 (2020).

29. M. G. Palmgren, P. Nissen, P-type ATPases. Annu Rev Biophys 40, 243–266 (2011).

30. R. W. Albers, Biochemical aspects of active transport. Annu Rev Biochem 36, 727–756 (1967).

31. R. L. Post, S. Kume, T. Tobin, B. Orcutt, A. K. Sen, Flexibility of an active center in sodium-plus-potassium adenosine triphosphatase. J Gen Physiol 54, 306–326 (1969).

32. M. Timcenko et al., Structure and autoregulation of a P4-ATPase lipid flippase. Nature 571, 366–370 (2019).

33. A. Davit-Spraul et al., ATP8B1 and ABCB11 analysis in 62 children with normal gamma-glutamyl transferase progressive familial intrahepatic cholestasis (PFIC): phenotypic differences between PFIC1 and PFIC2 and natural history. Hepatology 51, 1645–1655 (2010).

34. H. Egawa et al., Intractable diarrhea after liver transplantation for Byler’s disease: successful treatment with bile adsorptive resin. Liver Transpl 8, 714–716 (2002).

35. L. Bai et al., Autoinhibition and activation mechanisms of the eukaryotic lipid flippase Drs2p-Cdc50p. Nat Commun 10, 4142 (2019).

36. M. Chalat, K. Moleschi, R. S. Molday, C-terminus of the P4-ATPase ATP8A2 functions in protein folding and regulation of phospholipid flippase activity. Mol Biol Cell 28, 452–462 (2017).

37. S. H. W. Scheres, Amyloid structure determination in RELION-3.1. Acta Crystallogr D Struct Biol 76, 94–101 (2020).

38. S. Q. Zheng et al., MotionCor2: anisotropic correction of beam-induced motion for improved cryo-electron microscopy. Nat Methods 14, 331–332 (2017).

39. M. Su, goCTF: Geometrically optimized CTF determination for single-particle cryo-EM. J Struct Biol 205, 22–29 (2019).

40. P. B. Rosenthal, R. Henderson, Optimal determination of particle orientation, absolute hand, and contrast loss in single-particle electron cryomicroscopy. J Mol Biol 333, 721–745 (2003).

41. P. Emsley, B. Lohkamp, W. G. Scott, K. Cowtan, Features and development of Coot. Acta Crystallographica Section D 66, 486–501 (2010).

42. A. Waterhouse et al., SWISS-MODEL: homology modelling of protein structures and complexes. Nucleic Acids Res 46, W296–W303 (2018).

43. D. Liebschner et al., Macromolecular structure determination using X-rays, neutrons and electrons: recent developments in Phenix. Acta Crystallogr D Struct Biol 75, 861–877 (2019).

44. Schrodinger, LLC (2015) The PyMOL Molecular Graphics System, Version 1.8.

45. E. F. Pettersen et al., UCSF Chimera--a visualization system for exploratory research and analysis. J Comput Chem 25, 1605–1612 (2004).

