## Supplementary material for "Structural insights into the activation of autoinhibited human lipid flippase ATP8B1 upon substrate binding": Figures S1 to S10;Table S1;SI References


**This PDF file includes:**


Figures S1 to S10
Table S1
SI References

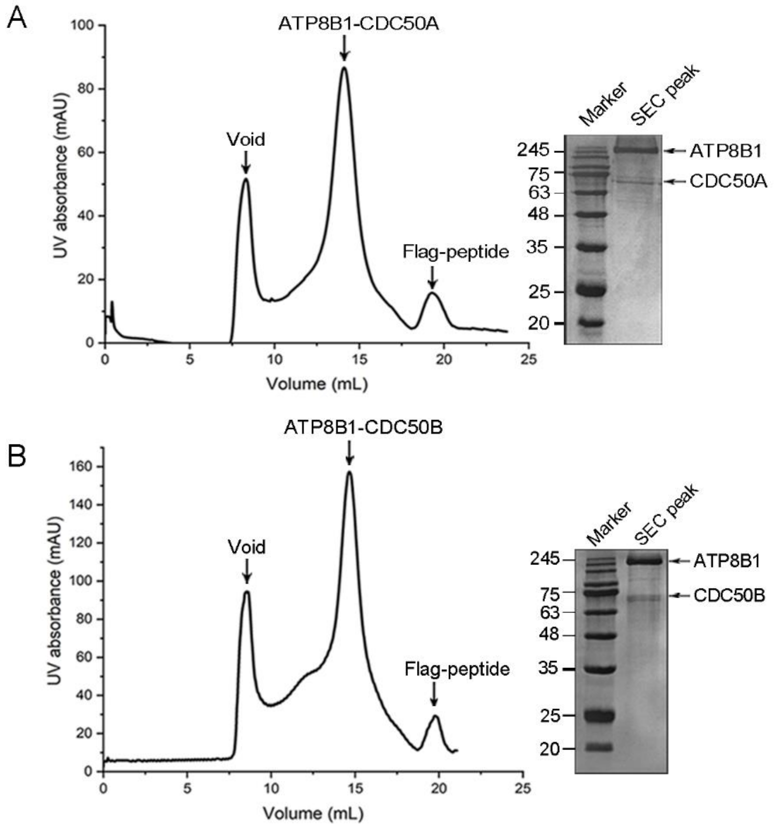


**Fig. S1.** Purification of ATP8B1-CDC50A and ATP8B1-CDC50B. Representative size-exclusion chromatography profiles and SDS-PAGE analysis of (A) ATP8B1-CDC50A and (B) ATP8B1-CDC50B.

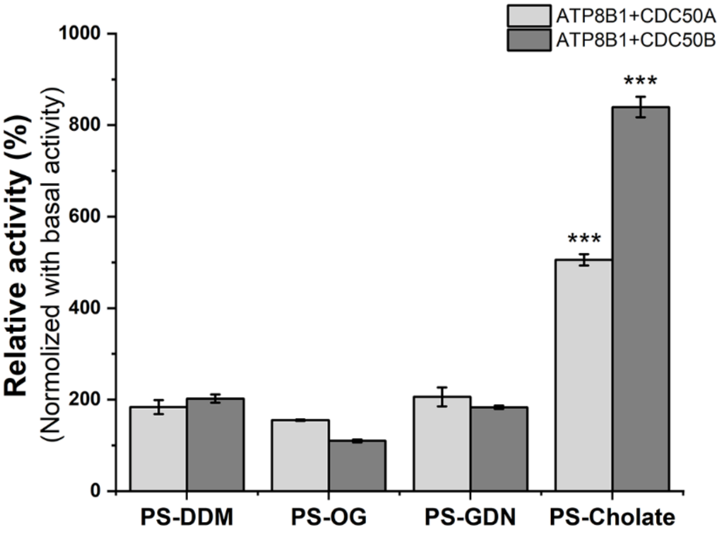


**Fig. S2.** The ATPase activity assays in the presence of PS solubilized in various detergents. Data were normalized against the basal activities of ATP8B1-CDC50A or ATP8B1-CDC50B, respectively. At least three independent assays were performed to calculate the means and standard deviations, and the data are presented as the means ± S.D. Two-tailed Student's t-test is used for the comparison of statistical significance. The p values of <0.05, 0.01 and 0.001 are indicated with *, ** and ***, respectively.

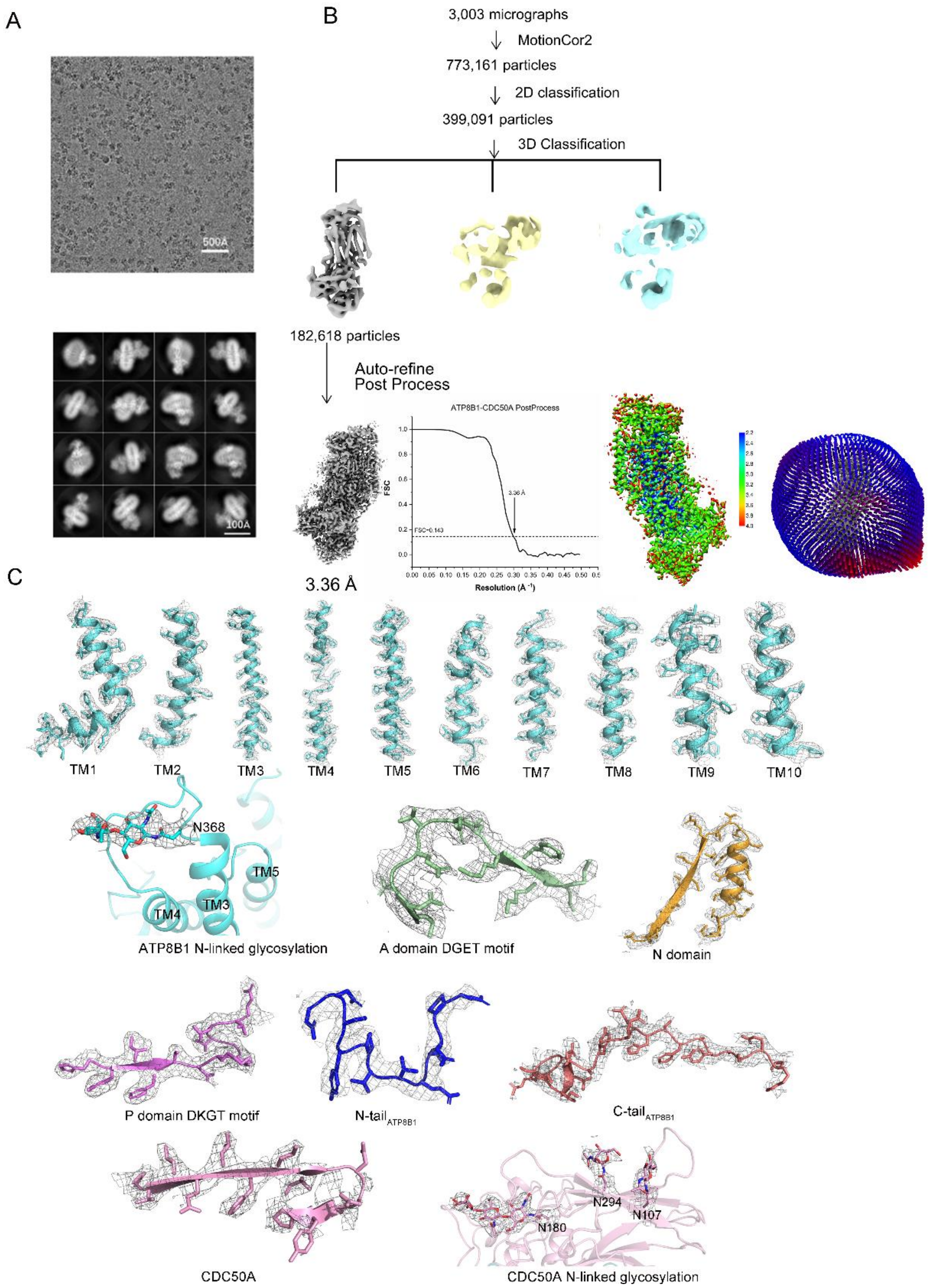


**Fig. S3.** Data processing and model building of the apo ATP8B1-CDC50A complex. (A) Representative cryo-EM image (upper) recorded on a 300 kV Titan Krios with a K2 camera and the 2D class averages of cryo-EM images (down) of ATP8B1-CDC50A. (B) Data processing workflow of the single particle image processing, the local resolution analysis and Cross-validation FSC curves for map-to-model fitting. Map resolution was estimated with the gold-standard Fourier shell correlation 0.143 criterion (1). Local resolutions were estimated using Resmap with RELION3.1 (2). (C) Atomic model of ATP8B1-CDC50A in the density maps. All maps were contoured at 4σ. Figures were prepared with Pymol (3) or Chimera (4).

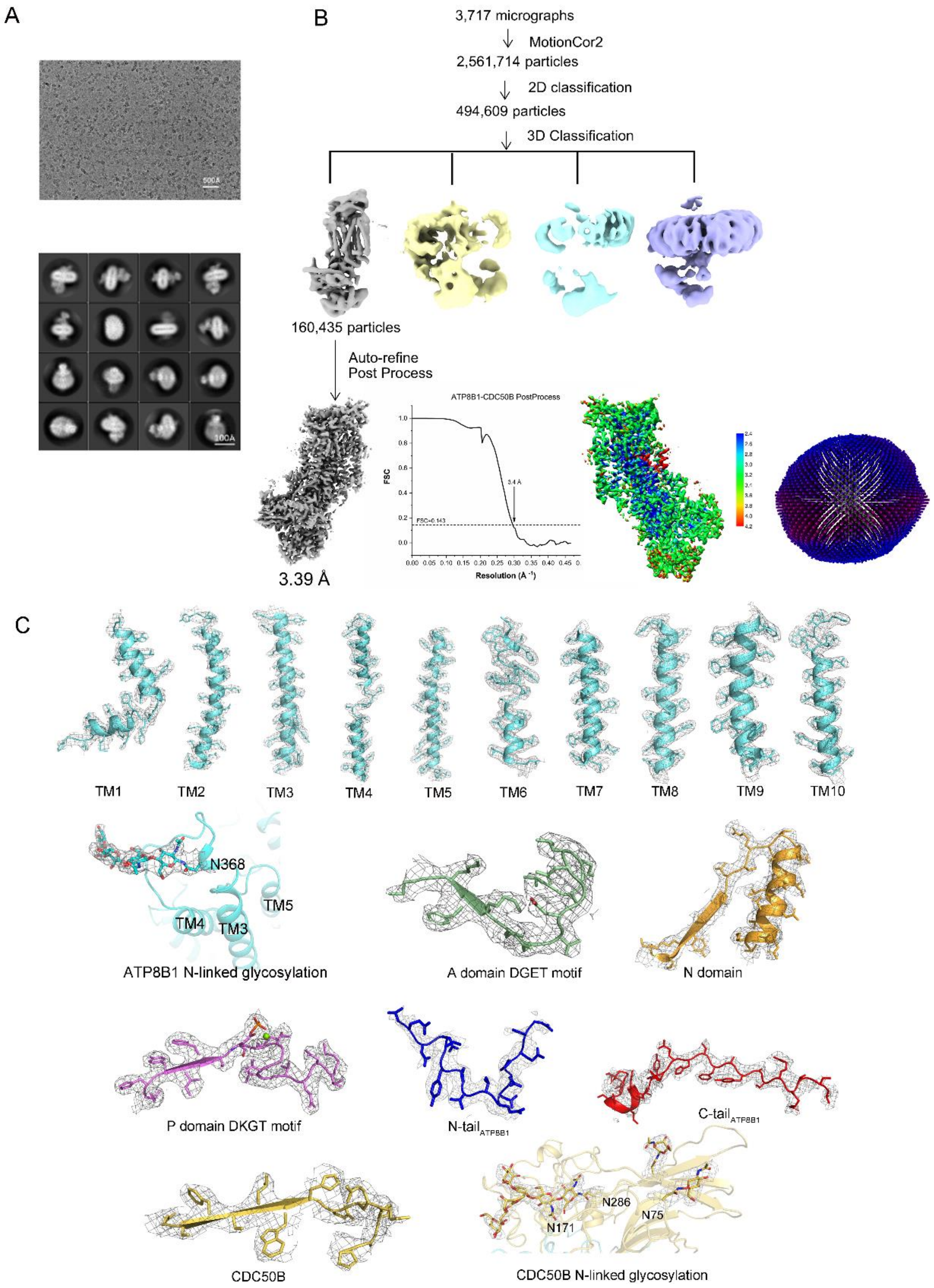


**Fig. S4.** Data processing and model building of the apo ATP8B1-CDC50B complex. (A) Representative cryo-EM image (upper) recorded on a 300 kV Titan Krios with a K3 camera and the 2D class averages of cryo-EM images (down) of ATP8B1-CDC50B. (B) Data processing workflow of the single particle image processing, the local resolution analysis and Cross-validation FSC curves for map-to-model fitting. Map resolution was estimated with the gold-standard Fourier shell correlation 0.143 criterion (1). Local resolutions were estimated using Resmap with RELION3.1 (2). (C) Atomic model of ATP8B1-CDC50B in the density maps. All maps were contoured at 4σ.

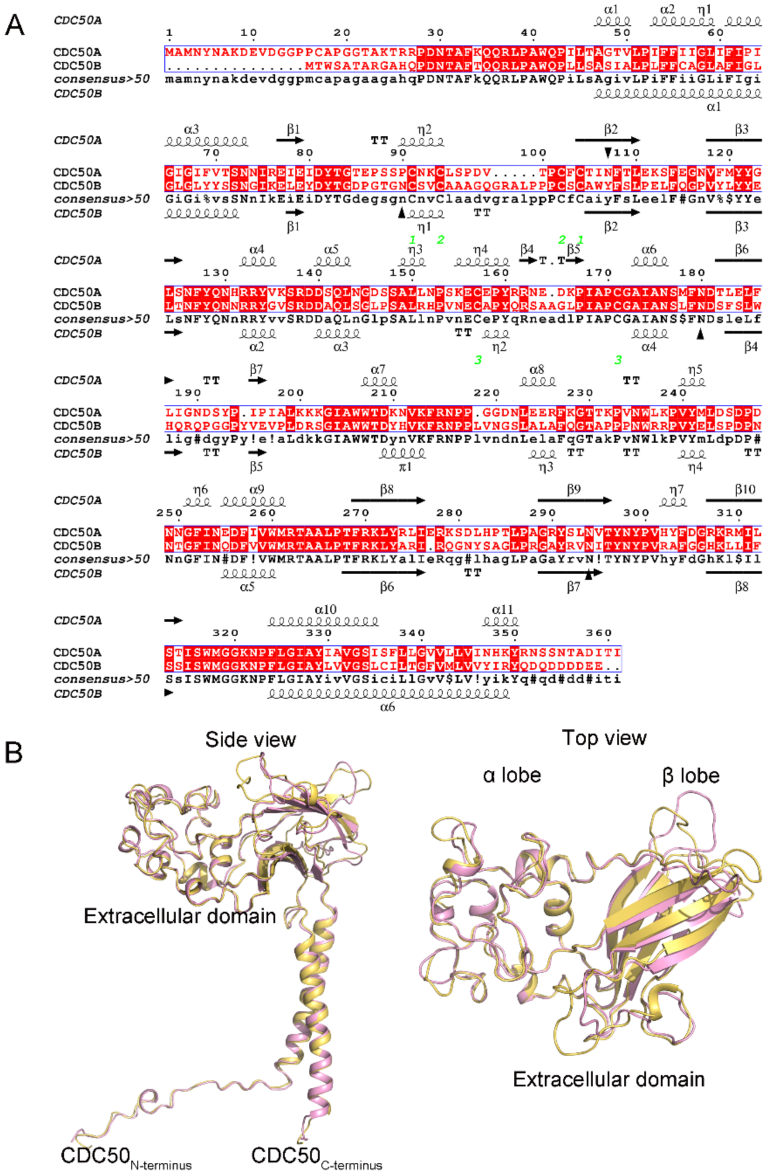


**Fig. S5.** Comparison of CDC50A and CDC50B. (A) Sequence alignment of human CDC50A and CDC50B. The secondary structure features deduced from CDC50A (upper) and CDC50B (down) are overlaid using ESPript 3.0 (5). (B)The side and top views of the superposition of overall CDC50A and CDC50B structures, colored in pink and yellow, respectively.

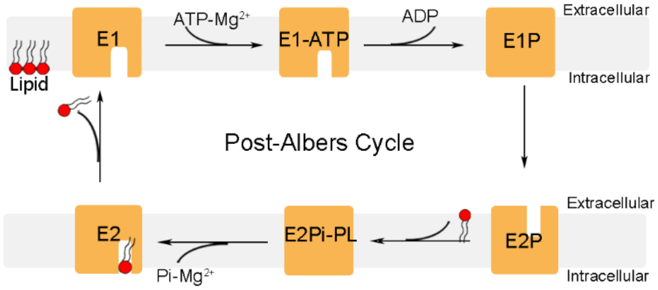


**Fig. S6.** Post–Albers cycle of P4-ATPases (6).

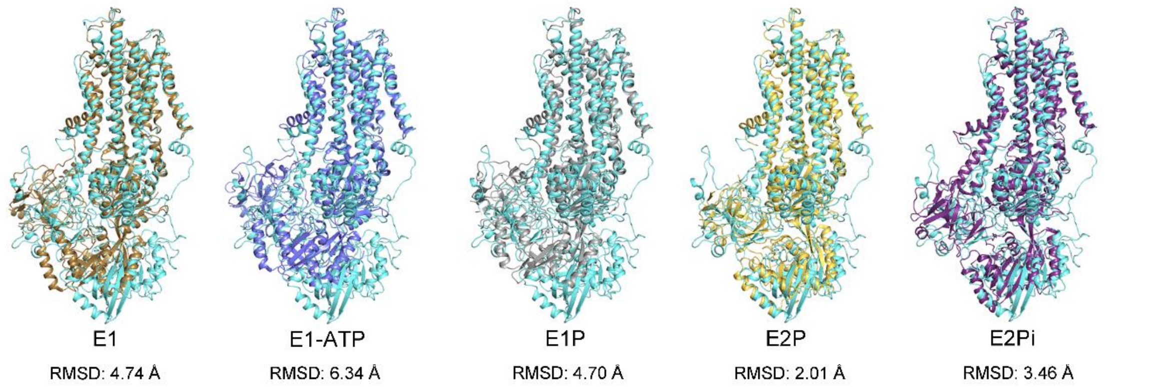


**Fig. S7.** Structure superpositions between ATP8B1 and ATP8A1 in different intermediate states. The apo-form ATP8B1 is colored in cyan, and ATP8A1 at various states are shown in different colors. The root-mean-square-deviation (RMSD) values to various intermediate states of ATP8A1 are shown at the bottom.

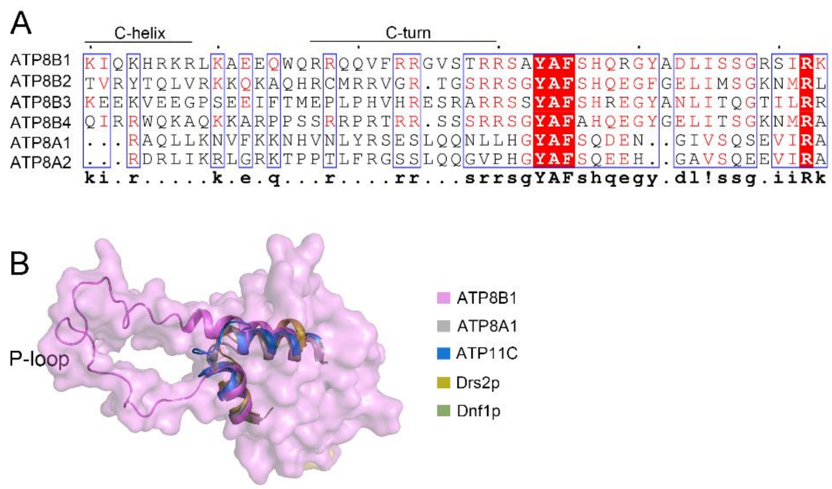


**Fig. S8.** Three positively charged regions. (A) Sequence alignment of C-tail of human P4-ATPase from Class 1, C-helix and C-turn of ATP8B1 are marked. (B) Structural comparison of the P domain. The P domain from ATP8B1 is shown as surface, and the P-loop shown as cartoon. The other P4-ATPases are shown as cartoon in different colors.

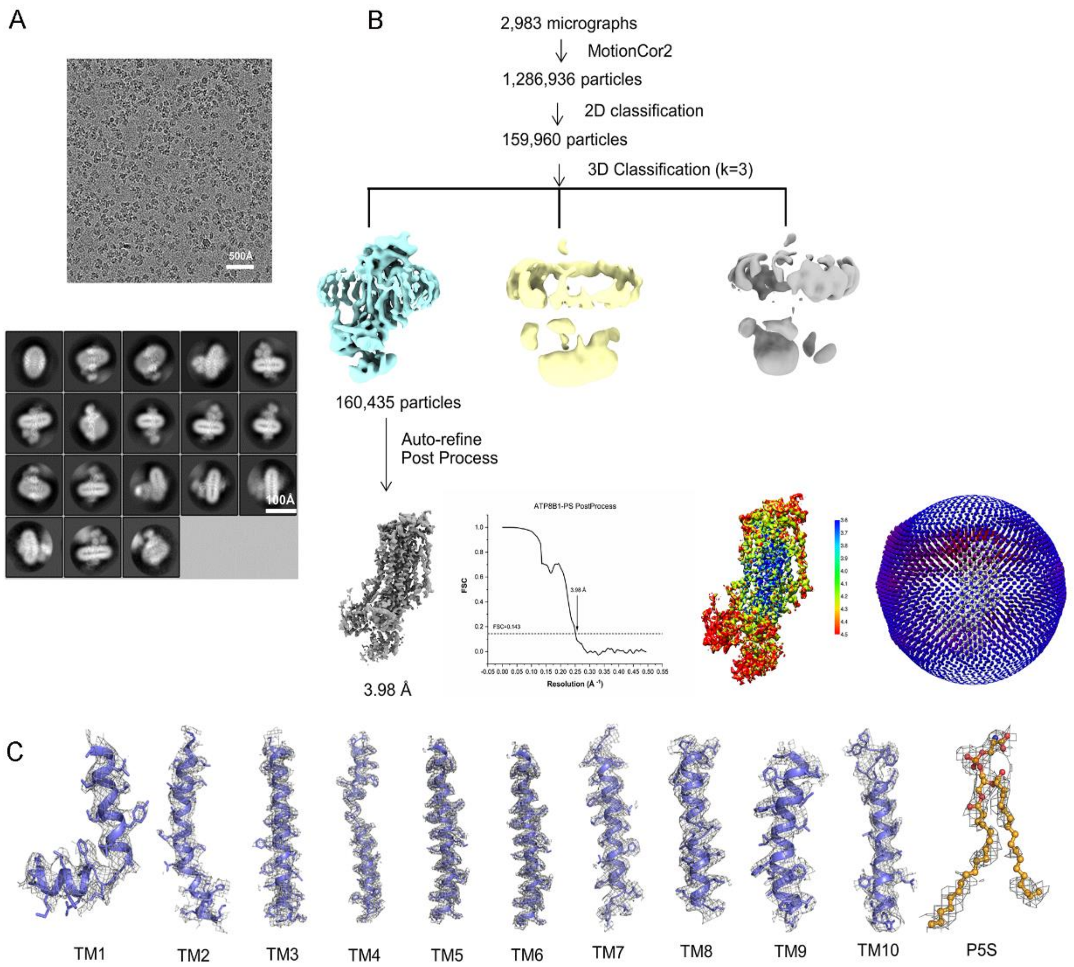


**Fig. S9.** Data processing of the PS-bound ATP8B1-CDC50A complex. (A) Representative cryo-EM image (upper) recorded on a 300 kV Titan Krios with a K2 camera and the 2D class averages of cryo-EM images (down) of PS-bound ATP8B1-CDC50A. (B) Data processing workflow of the single particle image processing, the local resolution analysis and Cross-validation FSC curves for map-to-model fitting. (C) The TMs and PS atomic model of ATP8B1-CDC50A in the density maps. The maps of TMs were contoured at 4σ, with the map of PS is at 3σ.

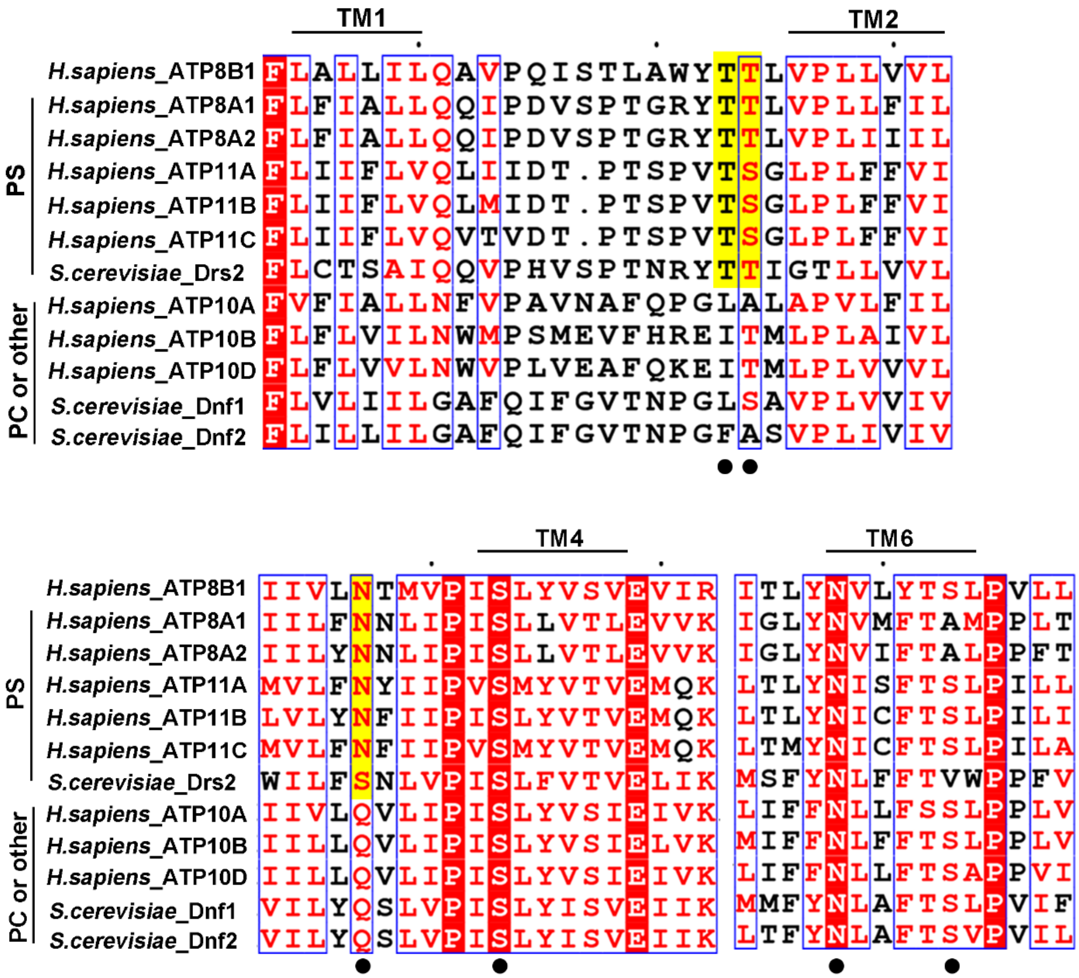


**Fig. S10.** Sequence alignment of the phospholipid-binding sites of P4-ATPases. PS transporters in top and that of PC in bottom. The substrate binding residues of ATP8B1 are labeled with black dots.

**Table S1.** Cryo-EM parameters, data collection and refinement statistics.

| | ATP8B1-CDC50A | ATP8B1-CDC50B | PS-bound ATP8B1-CDC50A |
|---|---|---|---|
| **Data collection and processing** | | | |
| Magnification | 29,000 | 22,500 | 29,000 |
| Voltage (kV) | 300 | 300 | 300 |
| Electron exposure (e – /Å$^2$) | 60 | 60 | 60 |
| Defocus range (μm) | -1.5 to -2.5 | -1.5 to -2.0 | -1.5 to -2.5 |
| Pixel size (Å) | 1.01 | 1.06 | 1.01 |
| Symmetry imposed | C1 | C1 | C1 |
| Initial particle images (no.) | 773,161 | 2,561,714 | 1,286,936 |
| Final particle images (no.) | 182,618 | 160,435 | 159,960 |
| Map resolution (Å) | 3.36 | 3.39 | 3.98 |
| FSC threshold | 0.143 | 0.143 | 0.143 |
| Map resolution range (Å) | 2.02-999 | 2.12-999 | 2.02-999 |
| **Refinement** | | | |
| Initial model used (PDB code) | 6K7L | | |
| Map sharpening B factor (Å$^2$) | -134.001 | -132.809 | -181.728 |
| Model resolution (Å) | 3.36 | 3.39 | 3.98 |
| FSC threshold | 0.143 | 0.143 | 0.143 |
| **Model composition** | | | |
| Non-hydrogen atoms | 12,197 | 12,405 | 11,460 |
| Protein residues | 1,493 | 1521 | 1,414 |
| Waters | 0 | 0 | 0 |
| **R.M.S. deviations** | | | |
| Bond lengths (Å) | 0.008 | 0.004 | 0.007 |
| Bond angles (°) | 1.156 | 0.709 | 1.085 |
| **Validation** | | | |
| MolProbity score | 2.14 | 2.13 | 3.23 |
| Clashscore | 9.74 | 9.61 | 43.72 |
| Poor rotamers (%) | 0.3 | 0.99 | 3.38 |
| **Ramachandran plot** | | | |
| Favored (%) | 86.64 | 87.04 | 82.15 |
| Allowed (%) | 13.09 | 12.83 | 17.07 |
| Disallowed (%) | 0.27 | 0.13 | 0.78 |

**SI References**


1. P. B. Rosenthal, R. Henderson, Optimal determination of particle orientation, absolute hand, and contrast loss in single-particle electron cryomicroscopy. *J Mol Biol* **333**, 721-745 (2003).
2. S. H. W. Scheres, Amyloid structure determination in RELION-3.1. *Acta Crystallogr D Struct Biol* **76**, 94-101 (2020).
3. Schrodinger, LLC (2015) The PyMOL Molecular Graphics System, Version 1.8.
4. E. F. Pettersen *et al.*, UCSF Chimera--a visualization system for exploratory research and analysis. *J Comput Chem* **25**, 1605-1612 (2004).
5. X. Robert, P. Gouet, Deciphering key features in protein structures with the new ENDscript server. *Nucleic Acids Res* **42**, W320-324 (2014).
6. J. A. Lyons, M. Timcenko, T. Dieudonne, G. Lenoir, P. Nissen, P4-ATPases: how an old dog learnt new tricks - structure and mechanism of lipid flippases. *Curr Opin Struct Biol* **63**, 65-73 (2020).
